# Information Theory as a consistent framework for quantification and classification of landscape patterns

**DOI:** 10.1101/383281

**Authors:** Jakub Nowosad, Tomasz F. Stepinski

## Abstract

**Context:** Quantitative grouping of similar landscape patterns is an important part of landscape ecology due to the relationship between a pattern and an underlying ecological process. One of the priorities in landscape ecology is a development of the theoretically consistent framework for quantifying, ordering and classifying landscape patterns.

**Objective:** To demonstrate that the Information Theory as applied to a bivariate random variable provides a consistent framework for quantifying, ordering, and classifying landscape patterns.

**Methods:** After presenting Information Theory in the context of landscapes, information-theoretical metrics were calculated for an exemplar set of landscapes embodying all feasible configurations of land cover patterns. Sequences and 2D parametrization of patterns in this set were performed to demonstrate the feasibility of Information Theory for the analysis of landscape patterns.

**Results:** Universal classification of landscape into pattern configuration types was achieved by transforming landscapes into a 2D space of weakly correlated information-theoretical metrics. An ordering of landscapes by any single metric cannot produce a sequence of continuously changing patterns. In real-life patterns, diversity induces complexity – increasingly diverse patterns are increasingly complex.

**Conclusions:** Information theory provides a consistent, theory-based framework for the analysis of landscape patterns. Information-theoretical parametrization of landscapes offers a method for their classification.

## 1 Introduction

There is a continuing interest in assessing a degree of similarity between landscape patterns, ordering landscapes by a property of interest, and landscape classification. This is because of a relationship between an area’s pattern composition and configuration and ecosystem characteristics such as vegetation diversity, animal distributions, and water quality within this area (Hunsaker and Levine, 1995; Fahrig and Nuttle, 2005; Klingbeil and Willig, 2009; Holzschuh et al., 2010; Fahrig et al., 2011; Carrara et al., 2015; Arroyo-Rodríguez et al., 2016; Duflot et al., 2017).

The main focus of research on quantitative assessment of landscape patterns has been a development and application of landscape indices (see statistics of topics published in Landscape Ecology collected by Wu (2013)). Landscape indices (LIs) are algorithms that quantify specific spatial characteristics of landscape patterns; a large number of LIs have been developed and collected (McGarigal et al., 2002). In principle, it should be possible to quantify the whole landscape using a collection of different LIs, which together characterize the entire pattern. Multi-indices description of landscape patterns has indeed been used (see, for example, Cain et al. (1997) or Long et al. (2010)) and continue to be used. The problem with such an approach is an uncertainty as to which subset of the large number of existing LIs to choose without introducing an undue bias toward some aspects of the pattern.

This problem can be partially addressed by performing the principal components analysis (Riitters et al., 1995; Cushman et al., 2008) on LIs calculated for all landscapes in the dataset and using vectors consisting of top principal components instead of indices themselves. Principal components suppose to represent the few latent variables, which, although not directly measurable, represent fundamental and independent elements of the pattern’s structure. Nowosad and Stepinski (2018) applied principal components analysis to over 100,000 landscape patterns taken from the European Space Agency (ESA) global land cover map (ESA, 2017). They found that the two top components, which together explained 70% of the variance in the dataset, were sufficient to parametrize all patterns in this dataset. It enables assessment of similarity between patterns and, thus, their classification.

Another way to achieve landscape classification is through clusterings of their patterns. Cardille and Lambois (2009) and Partington and Cardille (2013) clustered land cover patterns using the Euclidean distance between their principal components calculated from LIs. Niesterowicz and Stepinski (2013, 2016) clustered land cover patterns using the Jensen-Shannon Divergence between co-occurrence matrices representing the patterns. Both methods yielded reasonable results. This notwithstanding, there is a number of issues with using clustering to find landscape pattern types (LPTs). The number of clusters needs to be set a priori and mostly arbitrarily, and a within-cluster pattern variation is not well-controlled (Niesterowicz and Stepinski, 2017). LPTs are a posteriori interpretations of clusters, but clusters change from one dataset to another. This means that LPTs obtained via clustering are not universal and apply only to a dataset from which they were derived.

Wickham and Norton (1994) were the first to propose a classification of landscapes into universal LPTs. Their classification scheme divides pattern configurations into three classes: matrix, matrix and patch, and mosaic. Thresholds on minimum and maximum values of areas constituting matrix and patches determine a configuration type assigned to a given pattern. This classification was used for classifying land cover patterns across the conterminous United States (Riitters et al., 2000). In this paper, we are going to present a method of parameterizing landscape pattern configurations that leads to the universal classification of landscapes into landscape pattern configuration types (LPCTs).

A separate but related research track pertains to an ordering of landscapes. Ordering is arranging landscapes in a linear sequence according to an increasing value of a parameter. The expectation is that such sequence shows a continuous progression of the pattern’s character. Frequently, this sought after character is its complexity. In general, complexity is a concept defying a precise definition. For example, the Webster’s dictionary defines a complex object to be “an arrangement of parts, so intricate as to be hard to understand or deal with.” In the case of landscapes, their complexity is related to the intricacy of their patterns.

Recently, several works (Claramunt, 2012; Altieri et al., 2018; Wang and Zhao, 2018) proposed to order landscapes using a concept of spatial entropy. Spatial entropy is a modification of the Shannon entropy designed to measure spatial intricacy of a pattern. Boltzmann entropy is another concept aiming at ordering landscape patterns by their complexity. It is named after a physicist, Ludwig Boltzmann, who used it (Boltzmann, 1866) to show a relationship between thermodynamic entropy and the number of ways the atoms or molecules of a thermodynamic system can be arranged. In a context of landscape patterns, a macrostate is an overall configuration of a pattern (which can be measured using a single index) and a microstate is a specific assignment of categories to individual cells under the condition of fixed landscape composition.

Cushman (2016, 2018) proposed to measure a macrostate by the total edge (*TE*) of the landscape. Thus, *TE* corresponds to a “temperature” in the original Boltzmann entropy as applied to thermodynamics. It is easy to imagine that many different landscape microstates correspond to the same value of *TE*. The Boltzmann entropy of a given pattern, *S*, is a logarithm of the number of microstates having the value of *TE* as calculated for this pattern. Thus, a set of landscapes could be ordered by their *S* values. Gao et al. (2017) proposed a different approach to calculating S in the context of gradient instead of the mosaic model of the landscape.

Whether the proposed orderings of landscapes yield sequences that indeed reflect continuously increasing complexity of patterns remains to be determined. To make such a determination a representative set of real-life landscapes needs to be ordered and evaluated. Most of the evaluations done so far used simulated landscapes which lack the character and diversity of form found in real-life landscapes. Demonstrations of orderings on real-life landscapes (Wang and Zhao, 2018; Gao et al., 2017) used two few landscapes to make a judgment.

The above overview of different approaches to quantification, ordering, and classification of landscape patterns reveals a lack of consistent methodology. Different aspects of pattern analysis were addressed using different approaches, and those approaches, with the exception of the Boltzmann entropy, were not rooted in any theory. The principal objective of this paper is to demonstrate that the Information Theory (IT) (Shannon, 1948), as applied to a bivariate random variable representing a landscape, constitutes a consistent, theory-based quantitative methodology addressing all aspects of pattern analysis. Information-theoretical measures describe composition and configuration of landscape patterns, one-dimensional parametrizations of patterns using these measures correspond to orderings, and two-dimensional parametrizations correspond to classifications.

In the second section, we describe our methodology which is consistent with IT as applied to a bivariate random variable. Because our description is thorough and customized to the case of the mosaic model of a landscape, this section doubles as a guide for the use of IT of a bivariate random variable for applications in landscape ecology. In the third section, we describe our evaluation dataset of landscape patterns which has been carefully chosen to represent all major configurational types. In the fourth section, we show orderings of the evaluation set with respect to different IT metrics and compare them to orderings based on the two principal components (Nowosad and Stepinski, 2018) and to an ordering based on the Boltzmann entropy (Cushman, 2018). In the fifth section, we show a two-dimensional parametrization of landscape patterns and demonstrate that it provides a basis for classification of landscapes into universal LPCTs. Discussion and conclusions follow in the sixth section.

## 2 Methodology: Information Theory

Consider a mosaic model of landscape represented by a grid of cells with each cell assigned a categorical class label from the set {*c*_1_,…, *c_K_*} where *K* is the number of landscape classes. Our basic units of analysis are not single cells but pairs of ordered adjacent cells. A pair is regarded as a bivariate random variable (*x, y*) taking values (*c_i_*, *c_j_*), *i* = {1,…,*K*}, *j* = {1,…,*K*}, where *x* is a class of the focus cell and *y* is a class of an adjacent cell. Using adjacent cells is the simplest way to take into account spatial relations when analyzing a pattern. We start our analysis by calculating the co-occurrence matrix (Haralick et al., 1973) which tabulates frequencies of adjacencies between cells of different classes. The co-occurrence matrix can be thought of as a 2D histogram of cell pairs in a pattern; each bin of the histogram indicates the number of (*c_i_, c_j_*) pairs. The adjacency is defined by the rook’s rule (4-connectivity) and we distinguish between frequencies of (*c_i_, c_j_*) pairs and frequencies of (*c_i_, c_j_*) pairs. Using other definitions of adjacency and/or unordered pairs is also possible (Riitters et al., 1996).

Probabilities of (*x, y*) are given by a joint probability *p*(*x* = *c_i_*, *y* = *c_j_*) – a probability of the focus cell having a class *c_i_* and an adjacent cell having a class *c_j_*. We calculate the values of *p*(*x* = *c_i_*, *y* = *c_j_*) by dividing the co-occurrence matrix by the total number of pairs in the pattern. The informational content of bivariate random variable (*x, y*) is given by the IT concept of joint entropy which is computable directly from *p*(*x, y*),

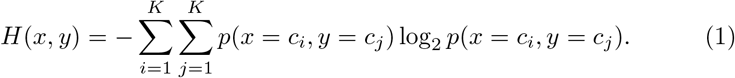

The value of *H*(*x, y*) is the number of bits needed on average to specify the value of a pair (*x, y*). It is also referred to as “an uncertainty.” We can interpret the uncertainty as the expected number of yes/no responses needed to determine a class of the focus cell and the class of the adjacent cell.

*H*(*x, y*) measures the diversity of heights of bins in a co-occurrence histogram. Recall that bins represent adjacencies, the larger the bin the more adjacencies of a corresponding type. If *H*(*x, y*) is small the histogram has few large bins – a landscape contains only a few types of adjacencies and thus its pattern is simple, If *H*(*x, y*) is large the histogram has many bins of similar height – a landscape contains many types of adjacencies and thus its pattern is complex. Thus, *H*(*x, y*) is a metric of an overall complexity of a pattern (see the *H*(*x, y*) ordering of the evaluation set of landscapes in Fig. 1).

**Fig. 1.**
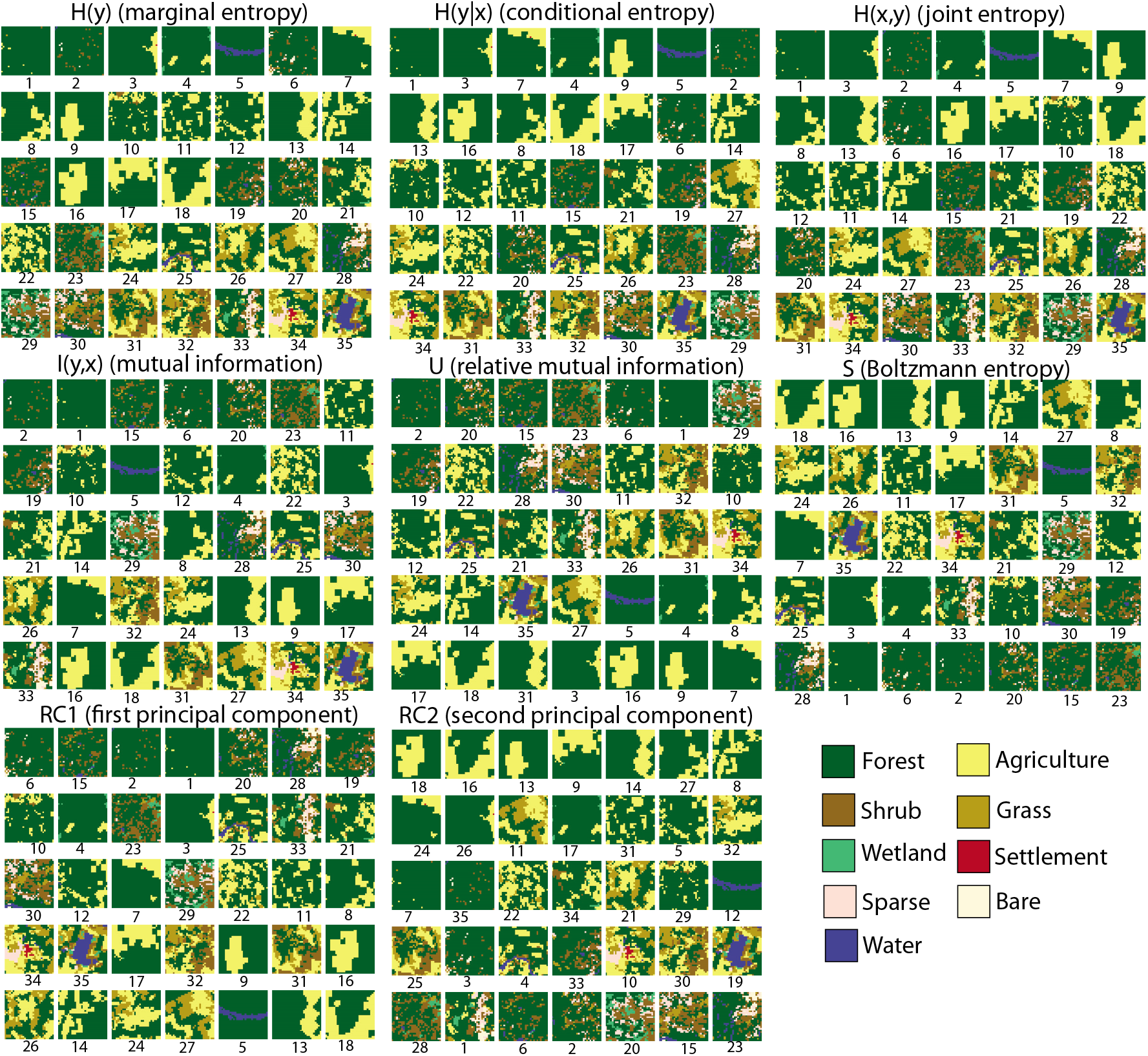
Linear orderings of evaluation landscape patterns by increasing value of indicated metrics. In each case, an ordering starts at the upper-left corner of a grid and proceeds row-wise. Numbers are the labels of patterns which are also ranks in *H*(*y*) ordering.

Next, we consider subsets of cell pairs such that a class of the focus cell is fixed. In such subset, the class of the adjacent cell is an univariate random variable *y*|*x* = *c_i_* taking values *y* = {*c*_1_,…,*c_K_*}. We can construct a 1D histogram, where bins correspond to frequencies of classes of adjacent cells in such subset. The variable *y*|*x* = *c_i_* has a probability distribution *p*(*y*|*x* = *c_i_*). The entropy of this distribution is 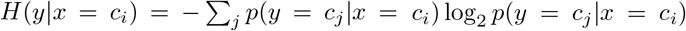. The value of *H*(*y*|*x* = *c_i_*) is the amount of bits needed on average to specify a class of an adjacent cell if the class of the focus cell is *c_i_*. It is also a diversity of adjacencies with class *c_i_*. If the value of *H*(*y*|*x* = *c_i_*) is small, cells of class *c_i_* are adjacent predominantly to only one class of cells, but if the value of *H*(*y*|*x* = *c_i_*) is large, cells of class *c_i_* are adjacent to many cells of many different classes. To obtain the full account of distribution of adjacencies we use the IT concept of conditional entropy,

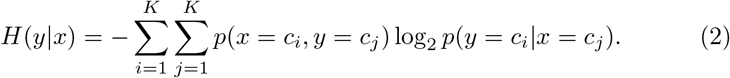

The conditional entropy, *H*(*y*|*x*) is an abundance-weighted average of values of *H*(*y*|*x* = *c_i_*) calculated for subsets of cells with different classes of the focus cell. *H*(*y*|*x*) is a metric of a configurational complexity of a pattern (see the *H*(*y*|*x*) ordering of the evaluation set of landscapes in Fig. 1). Note that the landscape with the highest configurational complexity is not the same as the landscape with the highest overall complexity because, even so it has a more intricate geometry it has fewer categories.

Finally, we consider a univariate variable *y* – a class of the adjacent cell in a pair of cells. Probability distribution of *p*(*y*) is obtained by marginalizing 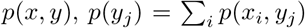. Informational content of *y* is computed using a standard Shannon entropy,

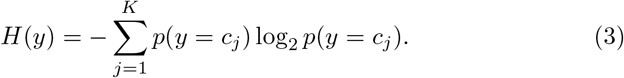

The value of *H*(*y*) is the number of bits needed on average to specify a class of cell. *H*(*y*) is a metric of a compositional complexity of a pattern, which is also frequently referred to as pattern diversity (see the *H*(*y*) ordering of the evaluation set of landscapes in Fig. 1).

We could also focus on variable *x* (a class of the focus cell) and calculate *H*(*x*). Because of the way the variables are defined, *H*(*x*) ≊ *H*(*y*), a small difference is due to a non-perfect symmetry of the co-occurrence matrix due to finite size of the landscape; for landscapes with a large number of cells the difference between *H*(*x*) and *H*(*y*) is negligible.

The IT chain rule formula (see, for example, Cover and Thomas (2012)) connects *H*(*x, y*), *H*(*y*|*x*), and *H*(*x*),

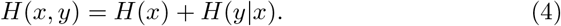

This formula shows that the informal statement – landscape patterns are characterized by both their composition and their configuration, which collectively define landscape structure – which is often found in landscape ecology papers, is not only a verbal description but has a quantitative justification.

One of the most useful concepts of IT is the mutual information, *I*(*y*, *x*), which quantifies the information that variable *y* provides about variable *x* (mutual information is symmetric so *I*(*x, y*) = *I*(*y, x*)). *I*(*y, x*) is given by the formula,

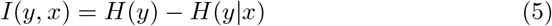

*I*(*y, x*) is a difference between uncertainty about the class of randomly drawn cell and a composition-weighted average uncertainty as to the class of the adjacent cell if drawn from subsets of pairs defined by a fixed value of the focus cell. It is also a difference between a diversity of cells’ categories and an average diversity of adjacencies (see the *I*(*y, x*) ordering of the evaluation set of landscapes in Fig. 1). The Jensen’s inequality (Jensen, 1906) assures that *I*(*y, x*) ≥ 0, so a diversity of adjacencies cannot exceed a diversity of categories.

Note that for real-life landscapes the value of *I*(*x, y*) tends to grow with a diversity of the landscape due to the spatial autocorrelation. The relative mutual information, *U* = *I*(*y, x*)/*H*(*y*), often referred to as an uncertainty coefficient, adjusts this tendency and has range between 0 and 1. It measures a difference between diversity of categories and diversity of adjacencies in terms of diversity of categories (see the *U* ordering of the evaluation set of landscapes in Fig. 1).

## 3 Evaluation dataset

In the introduction, we stressed the importance of using a complete dataset of real-life landscapes for an evaluation purpose. Such dataset needs to contain all feasible types of landscape pattern configurations. Global land cover maps offer a large dataset which contains rich variety of land cover patterns. We use a dataset (Nowosad et al., 2019) containing over a 1,600,000 (9km × 9km) landscapes extracted worldwide from the 300m resolution ESA 2015 global land cover map (ESA, 2017). To make the landscape more lucid, we reclassified the ESA map from the original 22 classes to 9 classes as listed in the legend to Fig. 1.

For these landscapes, we computed a set of 17 configurational landscape metrics (see Table 1 in Nowosad and Stepinski (2018) for details). Next, we calculated values of the top two principal components, *RC*1 and *RC*2 using the model of Nowosad and Stepinski (2018). Using these principal components we grouped landscapes into 35 types of pattern configurations (irrespective of their thematic content). For our evaluation dataset, we chose one exemplar from each of the 35 types of pattern configurations. We select exemplars only from landscapes with forest as a dominant theme so they are easier to compare visually, however, our results are theme-independent. The evaluation dataset is configurationally complete, at least for land cover landscapes at mesoscale.

For each of the 35 landscapes we calculated values of *H*(*y*), *H*(*y|x*), *H*(*x, y*), *I*(*y, x*), and *U*. We also calculated the value of Boltzmann entropy, *S* using the formula given in Table 6 of Cushman (2018) paper, and the values of the two top principal components, *RC*1 and *RC*2.

**Table 1.** Spearman’s rank correlation coefficients between different orderings

| ordering | $H(y)$ | $H(y\vert x)$ | $H(x,y)$ | $I(y,x)$ | $U$ | $RC1$ | $RC2$ | $S$ |
| --- | --- | --- | --- | --- | --- | --- | --- | --- |
| $H(y)$ | 1 | 0.93 | 0.97 | 0.59 | -0.13 | 0.25 | 0.55 | -0.06 |
| $H(y\vert x)$ | 0.93 | 1 | 0.99 | 0.30 | -0.42 | 0.02 | 0.75 | 0.17 |
| $H(x,y)$ | 0.97 | 0.99 | 1 | 0.41 | -0.30 | 0.12 | 0.68 | 0.07 |
| $I(y,x)$ | 0.59 | 0.30 | 0.41 | 1 | 0.67 | 0.71 | -0.22 | -0.67 |
| $U$ | -0.13 | -0.42 | -0.30 | 0.67 | 1 | 0.68 | -0.72 | -0.77 |
| $RC1$ | 0.25 | 0.02 | 0.12 | 0.71 | 0.68 | 1 | -0.49 | -0.93 |
| $RC2$ | 0.55 | 0.75 | 0.68 | -0.22 | -0.72 | -0.49 | 1 | 0.70 |
| $S$ | -0.06 | 0.17 | 0.07 | -0.67 | -0.77 | -0.93 | 0.70 | 1 |

$H(y)$ - marginal entropy, $H(y|x)$ - conditional entropy, $I(y,x)$ - mutual information, $U$ - relative mutual information, $RC1$ - first principal component, $RC2$ - second principal component, $S$ - Boltzmann entropy

## 4 Orderings of landscape patterns

Fig. 1 depicts orderings of evaluation patterns with respect to different metrics as indicated. All orderings are in the increasing value of a metric. They start from the upper-left corner of the grid of patterns and progress row-wise. The rankings for the *H*(*y*) ordering double as pattern labels; they are used in remaining orderings for quicker identification.

Because each metric (with the exception of *RC*1 and *RC*2) has an interpretation, it is interesting to see whether these interpretations agree with visual inspection of orderings. *H*(*y*) is interpreted as a diversity of cell categories. Although a visual inspection of landscapes sequenced by *H*(*y*) seems to confirm the overall tendency of increased compositional diversity, it also makes it very clear that very different patterns may have very similar levels of diversity (see, for example, landscapes #15 and #16). *H*(*y|x*) is interpreted as a diversity of cell adjacencies. Based on this interpretation, a sequence of landscapes ordered by *H*(*y|x*) should display increasingly heterogeneous (fine scale) patterns. Visual inspection shows that overall the heterogeneity of patterns in the sequence increases, but it also brings to our attention that landscapes with very similar values of *H*(*y|x*) are perceived as having very different heterogeneities (for example, see landscapes #5 and #2). Similar discrepancies can be observed in remaining orderings shown in Fig. 1.

These observations are explained by the fact that entropy is not an injective function of a histogram – different histograms may yield the same value of entropy. Thus, no linear ordering, based on entropy measure (note that *RC*1 and *RC*2 are indirectly also based, to some degree, on entropy-based indices) cannot be expected to produce a sequence with continuously changing character of pattern configuration, even if they show an overall trend in accordance with their interpretations. This brings into question practical values of complexity metrics such as the spatial entropy or the Boltzmann entropy. It is not that there is something wrong with these metrics, rather that they measure the same values for (sometimes) strikingly different patterns.

Table 1 lists rank correlations for orderings shown in Fig. 1. Values in this table confirm what is observed in Fig. 1, orderings of *H*(*y*), *H*(*y|x*), and *H*(*x, y*) are strongly correlated. Thus, in real-life landscapes diversity induces complexity. Because landscapes chosen for evaluation represent all feasible land cover pattern configurations, we expect that this observation extends to all land cover patterns. Thus, if a land cover pattern is diverse it is also complex. In fact, linear dependence between landscape complexity and its diversity has been observed in patterns present in Landsat images representing major Canadian ecoregions (Proulx and Fahrig, 2010).

The two mutual information metrics, *I*(*y, x*) and, especially, *U*, are poorly correlated with metrics *H*(*y*), *H*(*y|x*), and *H*(*x, y*), suggesting that mutual information provides a mostly independent, additional channel of information about a pattern. The first principal component, *RC*1 is moderately correlated with the mutual information and the second principal component, *RC*2, is moderately correlated with *H*(*y*), *H*(*y|x*), and *H*(*x, y*). Boltzmann entropy is moderately inversely correlated with the mutual information, strongly inversely correlated with *RC*1, and moderately correlated with *RC*2.

## 5 Landscape pattern configuration types

Rank correlations in Table 1 suggest using *H*(*y*) and *U* as the two parameters to utilize in a 2D parametrization of landscape patterns because they are the least correlated of all information-theoretical metrics. Fig. 2A is a graph showing such parametrization, hereafter we refer to it as the *HYU* diagram for short. Diagrams (not shown here) which use *H*(*y|x*) or *H*(*x, y*) instead of *H*(*y*) differ in details from the *HYU* diagram but have a similar overall character.

**Fig. 2.**
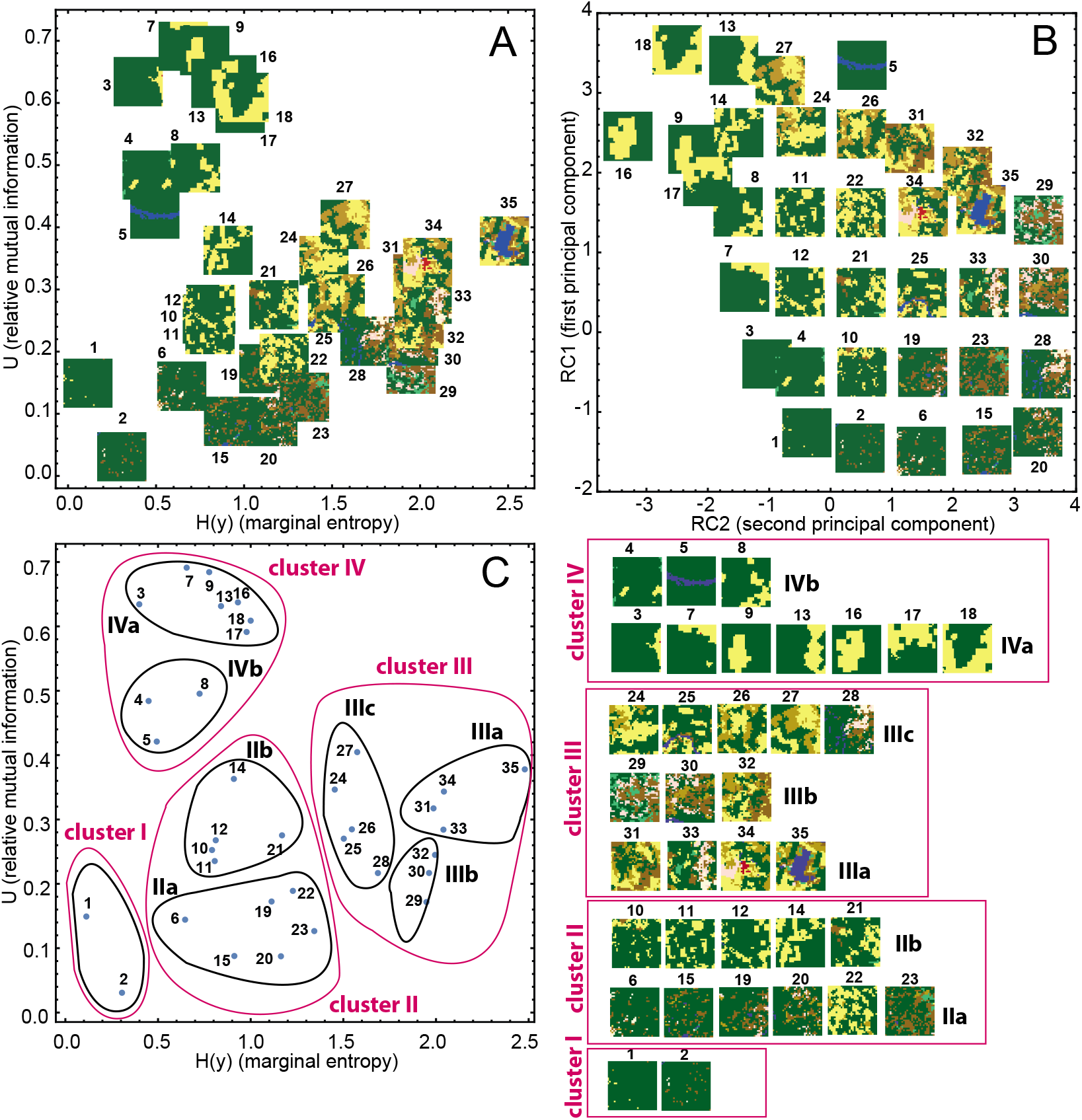
(A) Organization of evaluation landscape patterns by *H*(*y*) and *U*. (B) Organization of evaluation landscape patterns by the top two principal components *RC*2 and *RC*1. Landscapes are marked using labels introduced in Fig. 1. (C) Hierarchical clustering of landscapes into four LPCTs (red-colored contours) and eight LPCTs (black-colored contours). (D) Depiction of landscapes assigned to different LPCTs.

By analyzing the *HYU* diagram (possibly referring to Fig. 2D for an unobscured view of landscape patterns if necessary) it can be verified that it organizes landscape patterns in such a way that patterns placed in nearby locations on the diagram have similar configurations, and patterns placed in distant locations of the diagram have different configurations. Thus, there is a continuous relation between location of points on the *H*(*y*) − *U* plane and configurations of landscapes represented by these points. Because our evaluation landscapes have been chosen to represent all feasible land cover pattern configuration types this desirable feature of the *HYU* diagram extends to all land cover patterns.

The reason why a 2D parametrization succeeds in grouping similar patterns while 1D parametrization doesn’t is the presence of additional information that brakes a degeneracy (many-to-one mapping) of entropy-based measures. For example, very different patterns #15 and #16 are mapped to very similar values of *H*(*y*), but additional information – *U* – makes a distinction between them possible.

LPCTs can be extracted from the *HYU* diagram by clustering landscapes using the Euclidean distance between points on the *HYU* diagram as a measure of dissimilarity between patterns. Fig. 2C shows the result of hierarchical clustering (with Ward’s linkage) on 35 exemplar landscapes. Red-colored contours indicate clustering into four LPCTs and black-colored counters indicate clustering into eight LPCTs. Fig. 2D depicts landscapes belonging to individual clusters. It is clear from examining Fig. 2D that clusters group landscapes with similar configurations and thus can serve as LPCTs. Thematic content of landscapes within a single LPCT may differ as the *HYU* does not take it into consideration. To obtain a classification based on configuration and thematic content, LPCTs need to be further classified with respect to their themes.

Fig. 2B shows the *RC*2–*RC*1 diagram, which is an empirical counterpart of the *HYU* diagram. In the majority of cases, patterns placed in nearby locations on the *RC*2–*RC*1 diagram have similar patterns, but the relationship between pattern similarity and landscapes closeness in the 2D plane is not as good as in the *HYU* diagram. Also, the logic of the organization of pattern placements on the *RC*2–*RC*1 diagram is different from the logic of pattern placements on the *HYU* diagram. Most importantly, the *RC*2 – *RC*1 diagram is not universal. Landscapes patterns coming from a dataset other than the nine-classes ESA 2015 map cannot be placed on this diagram, because the principal components model used to construct this diagram does not apply to them. For this reason, the IT-based *HYU* diagram is a better classification tool than the empirically-based *RC*2 – *RC*1 diagram.

## 6 Discussion and conclusions

This paper makes several contributions to the theory of quantification of landscape patterns. The major contributions are as follows. (a) Demonstrating that fundamental properties of landscape patterns can be quantified within the framework of the Information Theory as applied to a bivariate random variable. (b) Showing that ordering landscapes by values of a single metric cannot yield a sequence of continuously changing patterns. (c) Observing that in real-life land cover landscapes diversity induces complexity; pattern’s configurational complexity is proportional to the pattern’s diversity. (d) Finding a 2D parametrization of landscape configurations based on two weakly correlated IT metrics that groups similar patterns into distinct regions of the parameters space thus providing the basis for classification of landscapes into LPCTs.

The first contribution is of conceptual nature. We demonstrated that landscape patterns can be quantified by calculating the distribution of information in a bivariate variable which describes a pattern. This is conceptually different from using ad hoc landscape indices. Note that IT of bivariate random variable provides information about composition (diversity) and configuration (adjacencies), thus providing all fundamental information about landscape configuration that is needed (Riitters, 2018). From equation 5 and the Jensen’s inequality, it follows that *H*(*y*) ≥ *H*(*y|x*) or that composition is a dominant property of the pattern. Also, we found that, at least for landscapes in our evaluation set, configuration follows composition. Together, these results are almost identical to conclusions recently reached by Riitters (2018) on the basis of long experience in working with landscape patterns.

Could we come up with our parametrization by just using landscape indices? Yes, but only in the retrospect. Presented parametrization emerges naturally from the IT-based analysis. Once emerged it can be expressed in terms of landscape indices. *H*(*y*) is equivalent to the Shannon’s diversity index (SHDI) and *H*(*x, y*) is inversely proportional to the contagion index, contagion = 1 − *H*(*x, y*)/max[H(x, y)], (O?Neill et al., 1988; Li and Reynolds, 1993). From those correspondences and using equations 4 and 5, it follows that *I*(*y, x*) can be expressed as a linear function of the contagion and the SHDI.

Using IT for quantification of ecologically relevant patterns was proposed before (Proulx and Parrott, 2008; Parrott, 2010) but only in the context of measuring the complexity of ecological systems, that is, in terms of our nomenclature, in the context of linear ordering. Another distinctive feature of the present paper is a thorough explanation of IT concepts in the context of landscape ecology, which can serve as a guide for future applications.

The second contribution is important because it brings into question whether orderings of landscape patterns (for example, by values of their complexity) are useful. For such ordering to be useful ordered landscapes should display a continuously changing pattern. Our results (see Fig. 1) shows that this is not the case for *H*(*x, y*) and *H*(*y|x*), the two IT-based measures of complexity and also not the case for the Boltzmann entropy. We suggested a simple explanation of why this must be so. This shortcoming of orderings has not been noticed before because proposed orderings were tested on either synthetic patterns or on small and incomplete samples of real-life patterns (Wang and Zhao, 2018; Cushman, 2018). In contrast, we used an evaluation set of landscapes that includes all types land cover configurations.

Because *H*(*x, y*) is inversely proportional to the contagion index, the ordering of landscape patterns by values of *H*(*x, y*) (see Fig. 1) demonstrates that the contagion index, which is considered to be a measure of clumpiness, is not really a good indicator of this property. Although a deficiency of contagion index as a measure of landscape clumpiness has been previously pointed out by (Li and Reynolds, 1993; Riitters et al., 1996; He et al., 2000), here we demonstrate it clearly on real-life landscapes.

The third finding – landscape diversity induces landscape compositional complexity – agrees with intuition. A diversity of categories is a prerequisite of pattern intricacy. Moreover, we demonstrated that, in real-life landscapes, there is a high correlation between pattern’s diversity and its complexity. Diverse but geometrically simple landscapes are just not found in nature. The high correlation between *H*(*y*) and *H*(*y|x*) in real-life landscapes points to an additional interpretation of relative mutual information *U* and the *HYU* diagram. An equation *H*(*y|x*) = *αH*(*y*) + *δ* states that the complexity of a pattern is equal to its prediction from the linear model, *αH*(*y*), which reflects an observed correlation, plus a “residual” *δ*. Note that *α* ≤ 1 due to the Jensen’s inequality and that there is no intercept in the linear model because for the homogeneous landscape *H*(*y*) = *H*(*y|x*) = 0. Thus eq. 5 can be rewritten as

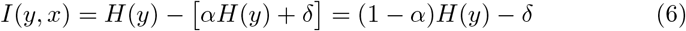

and the relative mutual information *U* can be expressed as

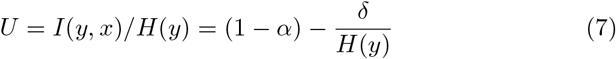

The term (1 − *α*) is an “expected” value of *U*, consistent with the observed correlation between composition and configuration. For our evaluation set of landscapes, this term is equal to 0.25. The second term is a part of *U* unaccounted for by the linear model. If a pattern is simpler than the linear model predicts *δ* is negative; such patterns are located above the *U* = 0.25 horizontal line on the *HYU* diagram. If a pattern is more complex than the linear model predicts *δ* is positive; such patterns are located below the *U* = 0. 25 horizontal line on the *HYU* diagram. Note that as the diversity of the composition increases the predictions of the linear model become more accurate.

Finally, our forth contribution has direct relevance to landscape classification into LPCTs. We have demonstrated that by using two weakly correlated IT metrics we can organize landscapes in such a way that landscapes with similar LPCTs are located in nearby locations on the 2D diagram. Thus, our *HYU* diagram is a de facto universal classifier of landscape pattern configuration types. It is an improvement over the classic method of Wickham and Norton (1994) inasmuch as it provides a more detailed classification of patterns’ configurations. However, our method does not consider thematic content of landscapes. For landscape classification based on configuration and thematic content, a post-processing step that further divides LPCTs on basis of themes is needed. This is a straightforward task, which, however, is beyond the scope of this paper. To facilitate classification of landscapes configurations via the *HYU* diagram we implemented *H*(*x, y*), *H*(*x*), *H*(*y|x*), and *I*(*y; x*) as the lsm_l_joinent, lsm_l_ent, lsm_l_condent, and lsm_l_mutinf functions in the R package landscapemetrics (Hesselbarth et al., 2019). The function accepts raster data as an input. Parameters include cells adjacency type (4-connected or 8-connected), and the type of pairs considered (ordered and unordered). Once these metrics are calculated for a set of landscapes, the *HYU* diagram can be constructed. Classification follows from a division of the *HYU* diagram by either manual or computational (clustering) means.

Since landscape patterns change with scale, future work will test the notion of the *HYU* diagram as the universal classifier on landscapes at different scales than in our present evaluation set. In particular, we plan on using land cover dataset having finer resolution than the ESA dataset, such as the National Land Cover dataset (NLCD, to test the *HYU* diagram on landscapes as small as 1km×1km. Because the *HYU* diagram is constructed on solid theoretical grounds, we expect that it would classify well landscapes at any scale.

## 7 Acknowledgments

This work was supported by the University of Cincinnati Space Exploration Institute.

